# Stress-adaptive biomaterials with tunable yielding architectures regulate organoid morphogenesis

**DOI:** 10.64898/2026.01.29.702625

**Authors:** James P.W. Reeves, Sabra Rostami, Mostafa Rammal, Andrei Bocan, Paula Lépine, Matthew J. Harrington, Thomas M. Durcan, Christopher Moraes

## Abstract

The yield stress at which biomaterials undergo plastic deformation limits the stresses that can be developed in encapsulated growing tissues. While mechanical properties of the matrix such as stiffness and viscoelasticity have a profound effect on cells, the role of yield stress has remained challenging to define. Here we design a self-healing granular hydrogel platform with supramolecular host-guest dynamic crosslinkers to precisely and quantitatively tune the stress at which the matrix repeatedly yields and reconfigures around tissues as they grow. Designed to provide similar mechanical constraints as a mesh stress ball, matrix yield stresses can be tuned between 12 and 370 Pa, while maintaining a storage modulus below ∼0.1kPa. We show that this range of yield stress is sufficient to promote or limit peripheral shedding in a model of non-adhesive cancer migration; and that early development of midbrain organoids is exquisitely sensitive to yield stress. Optimal yield stresses of only 25 Pa promoted budlike protrusions and large, lumenized neural rosettes, while variations as small as 10 Pa limited these phenotypes. These studies demonstrate that morphogenesis and tissue organization are exquisitely sensitive to yield stress, suggesting a new material property to target in designing biomaterials for disease modeling and regenerative medicine.

## 1. Introduction

Mechanical properties of the extracellular matrix (ECM) are now well-established to regulate cell fate and function, particularly in three-dimensional cultures^[1,2]^. Tunable, hypothesis-testing biomaterials have allowed precise identification of specific mechanical cues such as stiffness^[3]^, viscoelasticity^[4–6]^ and degradability^[7]^, and have provided design targets for applications in biomanufacturing^[8]^, therapeutic development^[9,10]^, and regenerative medicine^[11]^. Plasticity of the ECM, or the irreversible deformation that occurs under mechanical load is particularly important in directing tissue development^[12,13]^, as it provides an extracellular regulatory mechanism to release the stored deformation-induced mechanical energy generated during growth and morphogenesis^[14]^. Given that stress directly impacts biological activity, these extracellular mechanisms likely play an important role in differentiation^[15,16]^, morphogenesis^[17]^, and tissue destabilization^[18–20]^. However, developing biomaterials with defined plastic deformation profiles remains challenging.

Ubiquitous natural ECM such as collagen exhibit highly non-linear plastic deformation profiles^[21,22]^ which impact diverse remodeling processes in mammary branching^[23]^, hyperplasia, and fibrosis^[24]^. While natural ECM cannot be deterministically modified to tune the stresses supported by the material, synthetic alternatives can be designed to test these mechanobiological relationships. However, synthetic approaches are limited to either altering the stress-invariant time constants of plastic creep and stress relaxation^[25]^, or to manually manipulating local matrix properties via photochemistry^[7]^. Hence, it remains challenging to experimentally probe the causal relationship between ECM-mediated mechanical stress and biological function during plastic deformation, particularly for tissues that exhibit large morphogenetic transformations.

Here, we design a synthetic biomaterial that limits the stored mechanical energy in a tissue without time- or degradation-based dependencies, by locally yielding and reorganizing at precisely defined stresses. To achieve this, we leveraged a host-guest supramolecular dynamic crosslinking scheme, in which a ‘guest’ (adamantane; Ad) and ‘host’ moiety (β-cyclodextrin; β-CD) are allowed to temporarily and reversibly bind (Fig. 1a)^[26]^. These ‘ball-and-socket’ bonds can be mechanically detached, displaced, and rapidly reattached with neighbouring complementary moieties, creating a self-healing material^[27]^. Controlling the availability of binding sites should tune the yield stress of the polymer network (Fig. 1b). However, grafting guest and host moieties directly to hydrogel fibers that have some fraction of permanent crosslinks do not allow sufficient crosslinker mobility to rearrange after breakage^[27]^.

**Figure 1.**
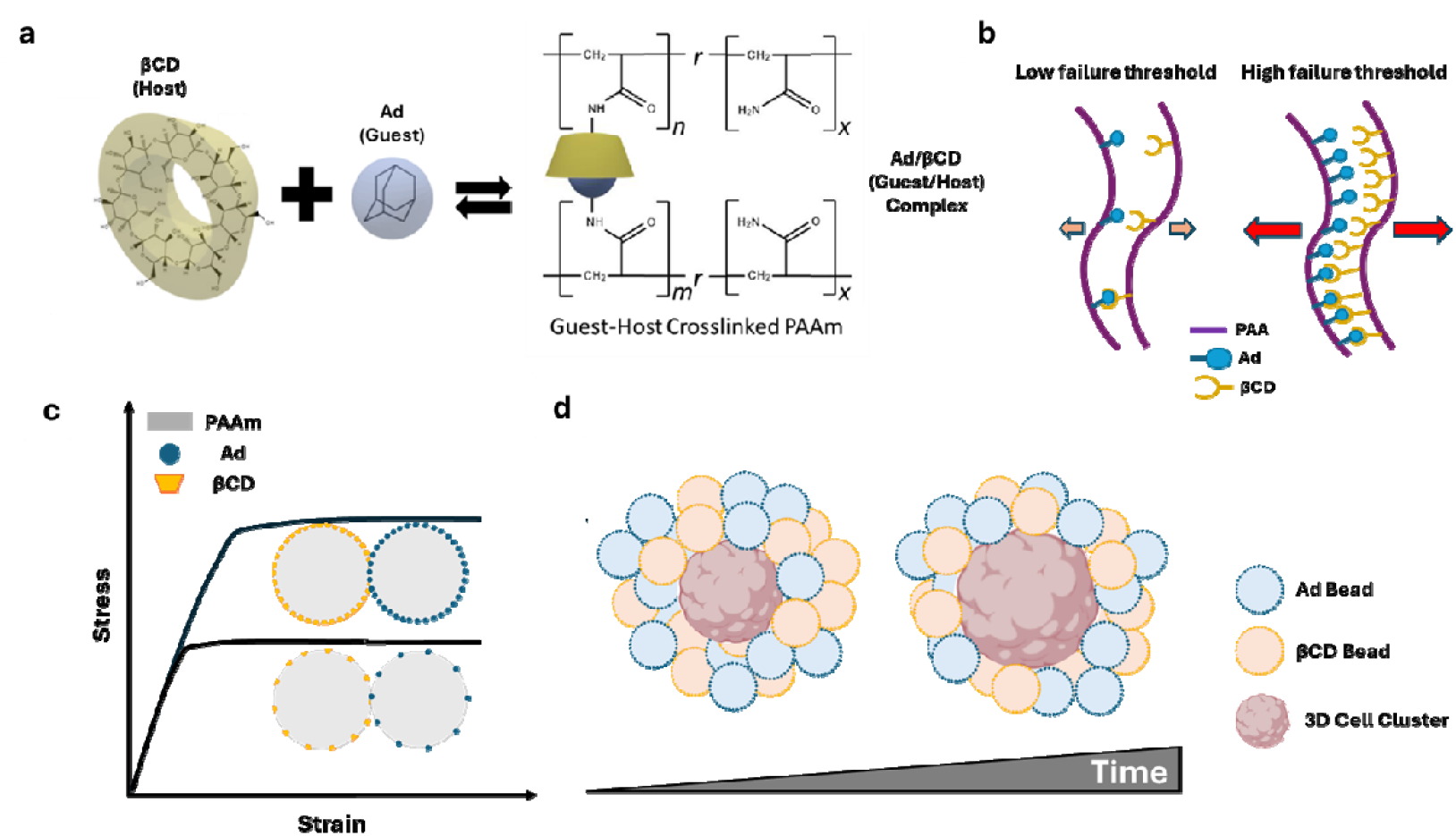
Design principle of yield-stress tunable granular gels. (a) Binding between the guest molecule, Adamantane (blue), to the host molecule, β-cyclodextrin (yellow), to form a dynamic crosslink between two polymeric surfaces. (b) Varying the quantity of guest/host pairs alters the stress required to detach surfaces. (c) Interacting complementary granules with varied guest host pairs can be used to regulate the yield stress required before the material undergoes plastic deformation. (d) Encapsulated tissues in the yield-stress tunable matrices can undergo growth and morphogenesis over time, while both generating and limiting the stresses they experience. Granules can locally break apart and rearrange themselves around the growing substrate to maintain an upper limit on local morphogenetic stresses.

To simultaneously obtain mechanical stability and crosslinker mobility, we separately grafted guest- and host-moieties to hydrogel microgranules (Fig. 1c) to obtain self-healing granular gels^[28]^. This concept has been previously utilized as a supporting matrix for 3D bioprinting^[29– 31]^ and as an injectable biomaterial^[32]^; and here we introduce control over the density of available crosslinking pairs to permit yielding and reorganization at quantitatively-defined stress thresholds (Fig. 1c). Tunability of dynamic adhesion between granules allows us to specify the yield point at which the matrix rearranges and reforms itself around the growing tissue (Fig. 1d). Much like a mesh stress ball that bulges when a mechanical stress is exceeded, this approach responds to regulate the local mechanical stresses generated within tissues during growth, by allowing stress-specific separation of the granules and local deconfinement of the tissue, without altering other critically important mechanical cues such as matrix stiffness.

## 2. Results

### 2.1. Surface functionalization of polyacrylamide granules can be tuned

Polyacrylamide was selected as the base material for the hydrogel granules used in this work, based on the materials’ well-established capacity to define and maintain linear elastic mechanical properties^[33]^, resist protein absorption^[34]^, accept surface modifications with specific extracellular matrix molecules ^[35]^ and form granules at scale via an oil-water emulsion process without alteration of mechanical properties^[36]^. We therefore characterized the parameter space achievable with this polymer. By edge-labelling the granules with fluorescent nanoparticles during gelation (red; Fig. 2a)^[37]^ granule size distribution was calculated as a function of vortex time. Polyacrylamide exhibited linear elastic behaviour up to 100% strain, with shear moduli between 50 Pa and 2.5 kPa, and negligible loss moduli (Supp. Fig. S1). Mean particle diameters can be tuned between ∼5 and ∼30 μm, with polydispersity indices ranging from 0.13 to 0.16, and no significant differences arising when incorporating βCD- or Ad host-guest linkers at their solubility limit (Fig. 2b). Longer vortex times produced smaller granules, but these could not be easily reclaimed using this fabrication method. While alternative strategies such as microfluidic or porous-glass droplet generation may produce more uniform granules^[38]^, polydispersity was desirable as it would enhance packing density and therefore increase the area for surface binding interactions between the granules.

**Figure 2.**
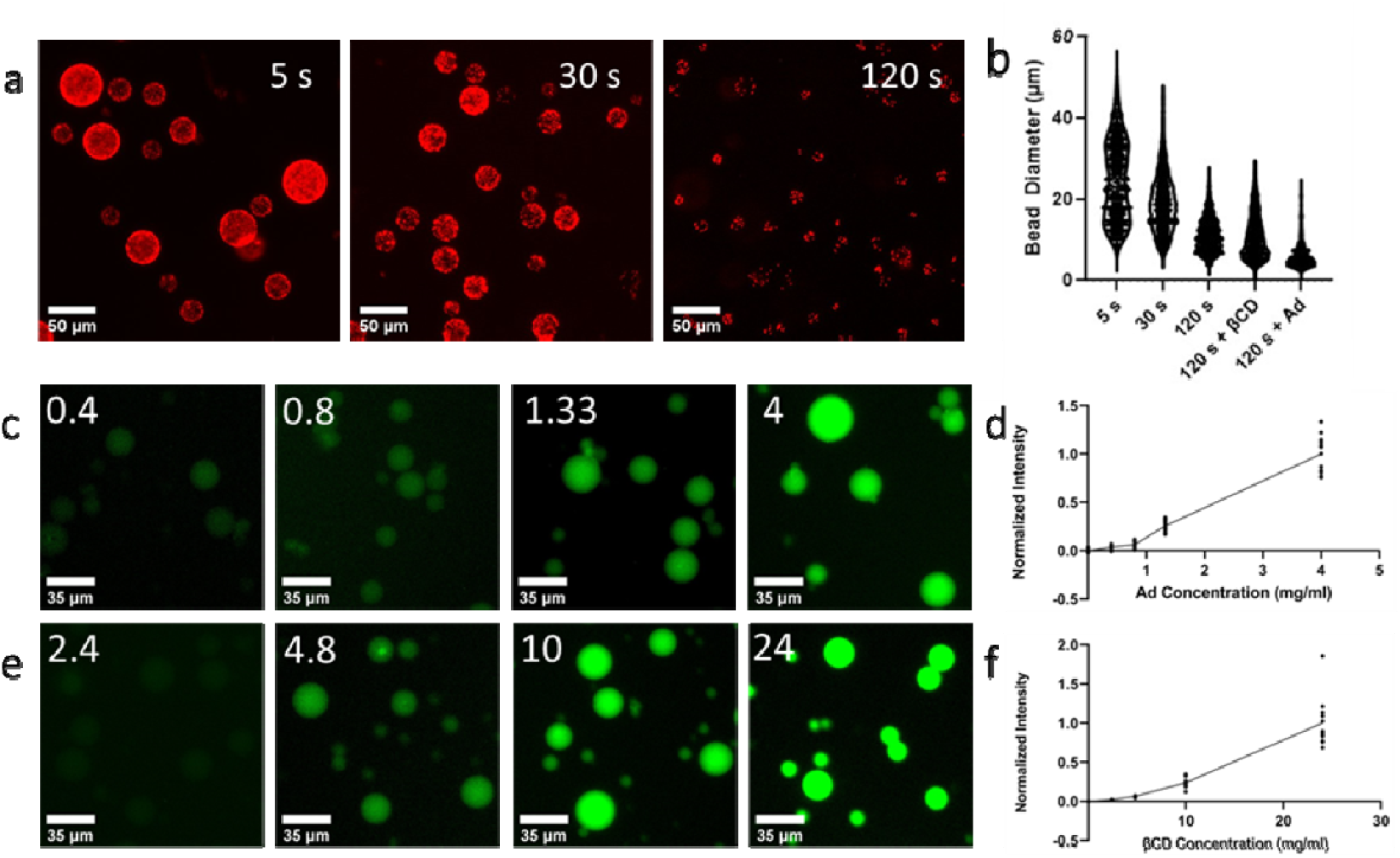
Characterization of individual granule properties. (**a)** Representative images of granules formed after 5 s, 30 s, and 120 s vortex times (red = edge-labelling fluorescent tracer nanoparticles). (**b)** Hydrogel granule size distributions for vortex times with and without βCD and Ad during fabrication (n>100). **(c-f)** Fluorescent images and normalized quantitative fluorescent intensities of **(c, d)** Guest (Ad-) and **(e, f)** host (βCD-) functionalized granules, based on brightness of a complementary fluorescent label (green).

We then verified that we could tune the concentration of supramolecular binding moieties in the granules using a fluorescent label for the guest- and host-moieties present after granule formation. As expected, fluorescent intensity increased with increasing crosslinker concentration (Fig. 2c-f; Supp. Fig. S2), determining the range of guest/host concentrations accessible with this approach.

### 2.2. Granule functionalization supports rapid and repeatable adhesion

By labelling the polyacrylamide material with fluorescein-methacrylate (green; β**-**CD granules) or embedded nanoparticle tracers (red; Ad granules) we could observe interactions between individual granules in a mixture using a standard fluorescent microscope. When mixed, only red- and green-labelled particles aggregated together, with softer granules visibly deforming upon adhesion to conform to the curved granule surfaces (Fig. 3a), while stiffer granules still bind but with smaller surface contact areas (Fig. 3b). No binding was observed between guest-guest (green-green) and host-host (red-red) particles.

**Figure 3.**
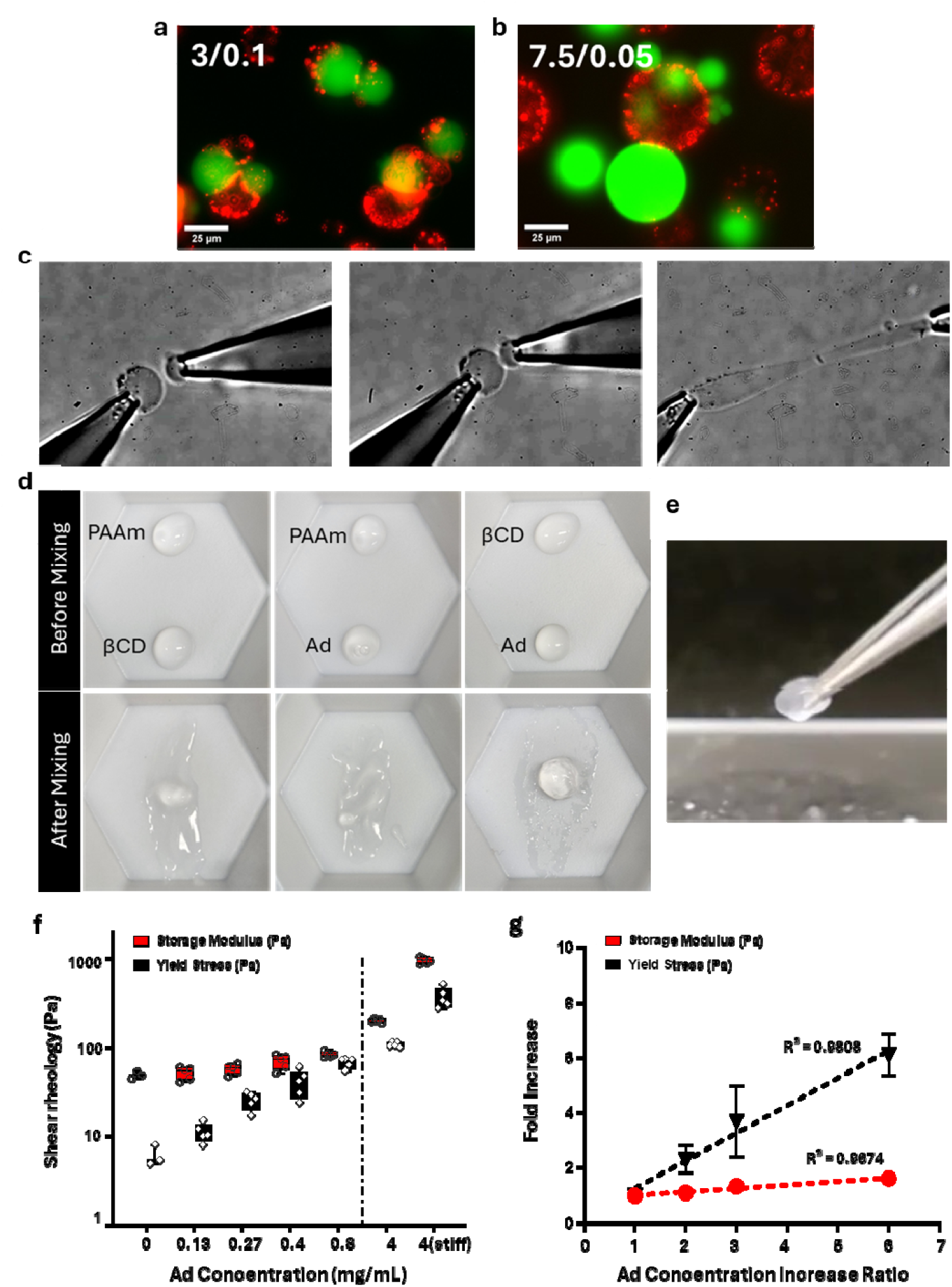
Guest- and host-decorated granule interactions. Granule-to-granule adhesion for **(a)** soft (G = ∼50 Pa) and **(b)** stiff (G = ∼2270 Pa) granules. **(c)** Representative images of contact and attempted separation of soft guest- and host-functionalized granules. (Scale bar = 20μm). **(d)** Mixing unfunctionalized/βCD-, unfunctionalized/Ad-, and βCD-/Ad-granule slurries. **(e)** Host-guest granule mixtures form a cohesive solid that can be picked up with tweezers. **(f)** Storage modulus and yield stress of mixed granular gels (1:1 mixing ratio of 24 mg/mL βCD-with varying concentration of Ad-granule slurries) **(g)** Fold increase of storage modulus and yield stress of the packed gel as a function of Ad concentration increase ratio at the 0.13 - 0.8 mg/mL range. Data presented as mean ± standard deviation, n = 4-6.

To determine whether binding times were appreciable, individual guest- and host-decorated granules were micromanipulated into contact. When unmodified polyacrylamide, guest-guest, or host-host granule pairs were touched and separated, they did not adhere (Supp. Fig. S3a, Supp. Movie SM1). In contrast, binding between guest- and host-decorated granules was instantaneous, and the soft granules withstood significant stretching before detachment (Fig. 3c; Supp. Movie SM2). To confirm that host-guest binding is repeatable for the same mating surfaces, we cyclically attached and detached an Ad-functionalized granule from a βCD-functionalized hydrogel at the highest crosslinker surface concentrations that we could synthesize (Supp. Fig. S3), and monitored granule deformation before detachment to confirm that surfaces adhered. Granules consistently stretched to an ellipsoidal shape while attached, and returned to a spherical shape after detachment, confirming that the granule is not mechanically damaged, even under the maximum stresses possible before adhesive failure. Furthermore, binding was repeatable between the same interacting surfaces over at least 10 attachment/detachment cycles (Supp. Fig. S3c) confirming that the supramolecular crosslinkers can support multiple detachment/re-attachment cycles.

### 2.3. Well-mixed granule slurries form cohesive matrices with tunable yield stresses

Diluted slurries of guest- and host-decorated granules (∼100 granules/μL) were then mixed, producing gels that were sufficiently cohesive to be manually picked up with tweezers (Figure 3d; 3e; Supp. Movie SM3). Mechanical vortexing and centrifugation produced more uniform mixing (Supp. Fig. S4) and was used in all subsequent experiments. By systematically modulating the mean granule size, stiffness, and functionalization, we measured the shear modulus and yield stress of the granular gels to determine how individual granule characteristics contribute to the global mechanical behaviour of the system.

We initially expected that decreasing the mean granule size would affect storage modulus and yield stress due to the increased interacting granule surface areas. However, no statistically significant differences in yield stress or storage modulus were observed for granules between 10 and 25 μm (Supp. Fig. S5). This is likely due to the polydispersity and relatively low stiffness of the granule populations tested, which would result in well-packed granular beds with maximal surface area interactions. Increasing the stiffness of the individual granules had a much more pronounced effect on mechanical properties of the granular gel (Supp. Fig. S6), likely due to the increased availability of host-guest binding sites that accompanies increased polymer content. Tuning this parameter produced materials with the highest stiffness and yield stress we have achieved in these experiments (G = 1 kPa, σ_y_= 400 Pa).

Reasoning that limiting either Ad- or βCD-surface concentrations would tune modulus and yield stress, we characterized the parameter space by tuning Ad-functionalization levels when mixed with the maximum concentration of βCD-coated granules (Fig. 3f, g) and vice versa (Supp. Fig. S7) with similar outcomes. Modifying the guest- or host-functionalization tuned matrix yield stress σ_y_ between 5 and 370 Pa (Fig. 3f). For all formulations with σ_y_ < 100 Pa the storage moduli were uniformly low and not statistically distinguishable (G < 0.1 kPa; Fig. 3g). Any minor variations in stiffness below this threshold are unlikely to significantly affect biological function^[39]^, and we therefore conclude that within this range, we can independently tune yield stress while maintaining a constant effective storage modulus.

### 2.4. Cancer cell invasive migration can be modulated by matrix yield stress

Cancer cells can make use of ECM adhesions to pull themselves away from the tumor body, but a growing body of evidence suggests that cancer cells can also invade into confined spaces using non-adhesive mechanisms^[40]^. In this first application to test the range and precision of the stress-yield granular gels, we sought to determine whether peripheral shedding via non-adhesive mechanisms can be regulated by the development of growth-induced mechanical stresses over time.

Aggregated T47D spheroids typically form tight boundaries and can escape as peripherally shed single cells^[41]^. We first confirmed that functionalized granules were not cytotoxic by screening against a T47D breast cancer cell line, and noted that cells spread, proliferated, and maintained viability over 3 days (Supp. Fig. S8). We next embedded T47D spheroids into a commercially available non-remodeling elastic polyvinyl-alcohol based gel, and into the granular gels developed in this work (Fig. 4). Although the light scattering properties of the granules makes live imaging challenging, live imaging is the only way to monitor peripheral shedding.

**Figure 4.**
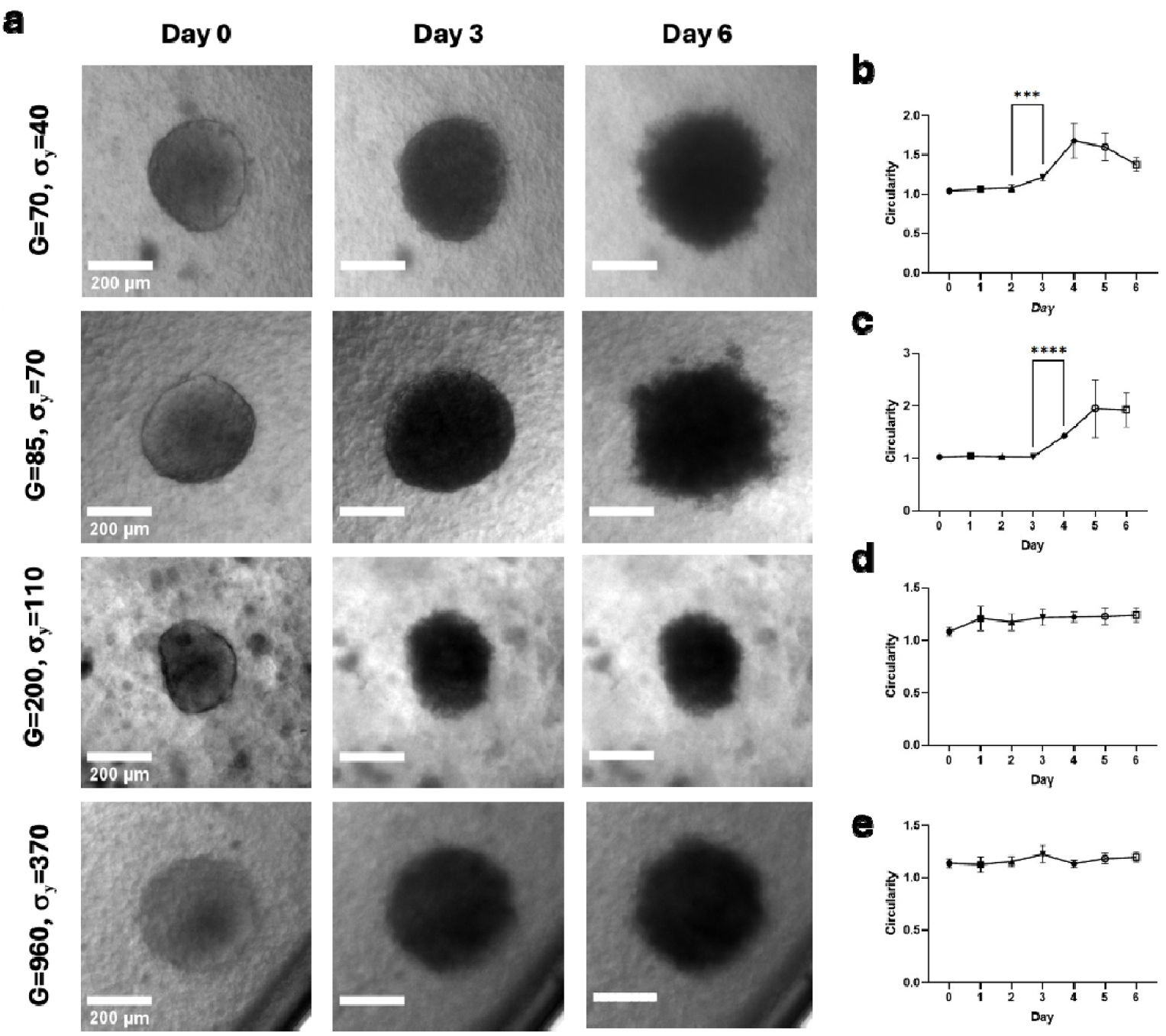
T47D aggregate growth in packed granular host-guest hydrogels. **(a)** Aggregates embedded in packed granular hydrogels of varying stiffness and yield points, at day 0, day 3, and day 6. G = storage modulus [Pa], σ_y_= yield stress [Pa]. **(b-e)** Invasiveness of the aggregate into the surrounding environment as measured by its changing circularity. (Data presented as mean ± standard deviation, n = 4-6, *** = p < 0.001, **** = p < 0.0001 via repeated measures ANOVA with Tukey post-hoc comparisons)

Over 6 days, spheroids in elastic gels as stiff as 1 kPa (Shear modulus, Supp. Fig. S9) and in yield-stress tunable granular gel formulations with comparable moduli (Fig 4) consistently grew larger, demonstrating that spheroid growth is not limited in the stiffness range presented by our granular gel systems (Supp. Fig. S10). We then quantified peripheral shedding by measuring spheroid circularity. As expected, no peripheral shedding was observed in elastic gels (Supp. Fig. S9), or in granular matrices with large yield stresses (σ_y_ ≥ 100 Pa; Fig 4d, e), demonstrating that the yield stress of the material can be specified to be sufficiently high to restrict shedding in this experimental model. Interestingly, however, peripheral shedding phenotypes were observed at earlier time points when the yield stresses are reduced (by Day 4 at σ_y_ = 70 Pa; and by Day 3 at σ_y_ = 40 Pa). This progression suggests that if internal stresses increase with time in culture, spheroids can gradually develop the stored mechanical energy required to shed cells via non-adhesion dependent mechanisms, suggesting that tumoral stress can develop over time, and is an important parameter in regulating peripheral shedding.

### 2.5. Early brain organoid architectures are exquisitely sensitive to matrix yield stress

We then asked whether matrix yield stress might impact internal tissue organization in a more complex model of tissue development, such as when an embryoid body undergoes growth, symmetry-breaking and organization during progression towards a midbrain organoid (MBO). Week-old embryoid bodies are typically grown in reconstituted basement matrix (rBM) such as Matrigel for several months, which is known to provide critically important chemical^[42]^ and mechanical ^[43]^ cues that regulate MBO architecture. Although the viscoelastic properties^[44]^, cell-mediated enzymatic degradation, matrix secretion, and mechanical non-linearities of rBM make it extremely challenging to determine the effective yield stress experienced at the organoid-matrix interface, matrix yielding must be required for organoids to grow and undergo morphogenesis, particularly during the first two weeks of culture when morphological changes are most pronounced.

To determine whether matrix yield stress is a critical regulatory parameter for early MBO morphogenesis, we cultured MBOs for two weeks in yield-stress tunable granular gels, while maintaining an rBM-like chemical stimulation profile using solubilized Matrigel (1.5%). Few dead cells were observed, indicating that the granular gels support sufficient delivery of nutrients and gas exchange (Fig. 5a). MBOs in conventionally-gelled rBM (Matrigel; 100%) displayed gross morphological bud-like protrusions (Supp. Fig. S11a). Similar bud-like protrusions were observed only at remarkably small yield stresses (12 ≤ σ_y_ ≤ 40 Pa; peak at 25 Pa; Fig. 5a), but MBOs in free media (σ_y_ = 0 Pa) or in gels with higher yield stresses (σ_y_ ≥ 70 Pa) were consistently rounded and smooth (quantitatively assessed in Fig. 5d). We next examined finer morphological features at the MBO edges, and noted that yield stresses ≥ 40 Pa induced a jagged ruffle-like phenotype (Fig. 5b), suggesting that when greater stresses are required to break through the surrounding matrix, stochastic redistribution of growth stresses would result in more focalized morphogenetic growth and expansion. These analyses demonstrate that gross morphogenetic processes are exquisitely sensitive to matrix yield stress, at resolutions of less than 10s of pascals.

**Figure 5.**
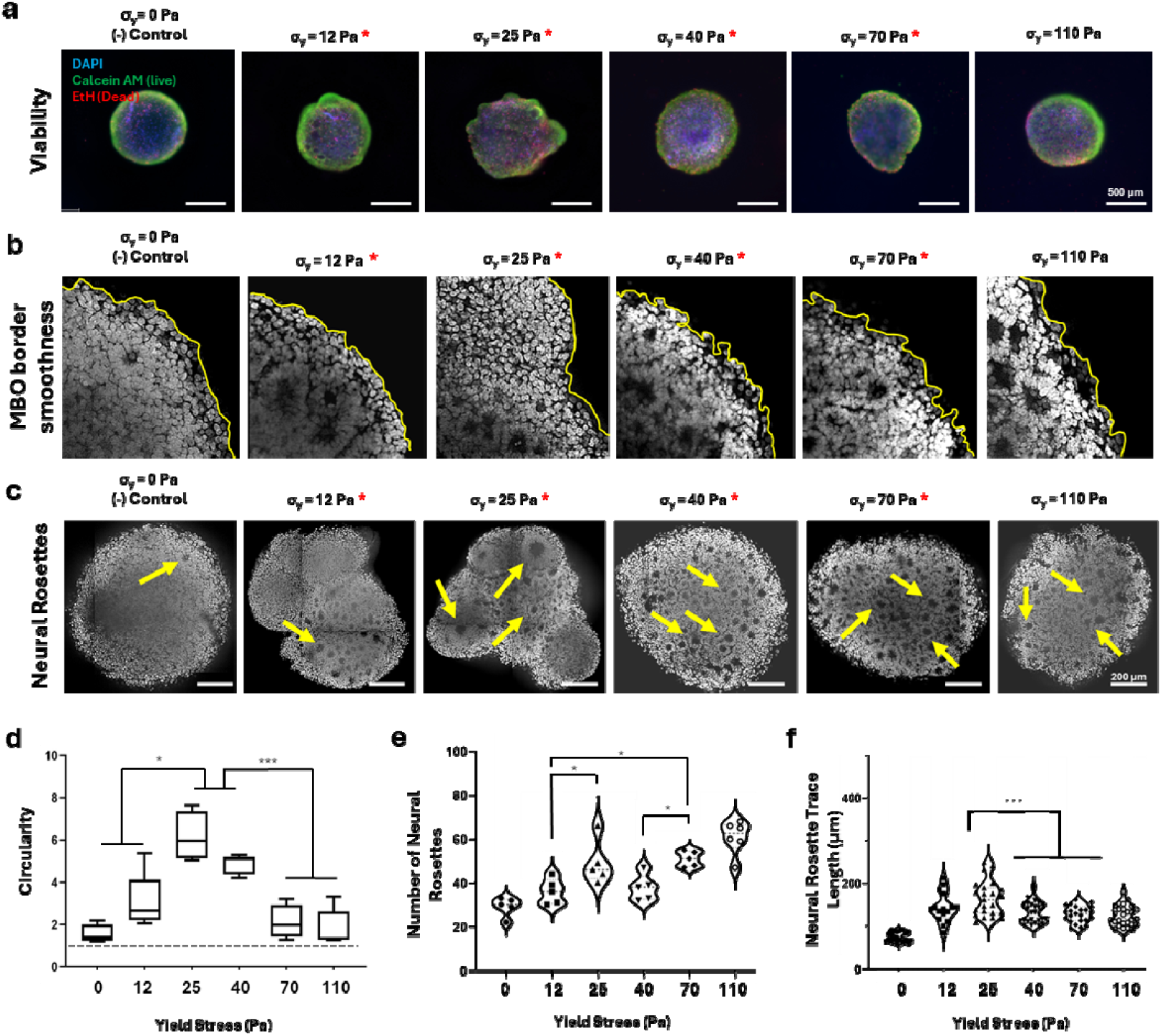
Midbrain organoids after two weeks in yield-stress tunable granular gels. Red stars indicate the conditions with statistically equivalent storage moduli and varying yield stress, σ_y_ = 0 denotes no gel condition; all cultures were supplemented with solubilized rBM to maintain chemical stimulation profile). **(a)** Gross overall MBO morphology captured via a projected stack of live/dead labelled images. **(b)** MBO periphery, yellow line denotes organoid edge. **(c)** Representative nuclei-labelled MBO sections; yellow arrows indicate example neural rosettes within the organoids. **(d-e)** Quantitative analysis of **(d)** circularity of the MBOs, **(e)** number of neural rosettes, and **(f)** trace length of the neural rosettes per organoid section. All data presented as violin or box plots denoting the mean and first to third quartile, and the whiskers span the range (n = 4-5 organoids; * p < 0.05, *** p < 0.001 by one-way ANOVA with Tukey post hoc comparisons).

Significant differentiation is not expected at this relatively early time point^[45]^. We confirmed this in conventional gelled Matrigel (100%) via confocal immunofluorescence for MAP2-labelled mature neurons and TH-labelled dopaminergic neurons at the organoid edges, and maintenance of Nestin, a marker for neural stem cells (Supp. Fig. S12). Interestingly however, expression of differentiation and stemness were all qualitatively improved within the organoid within the same optimal range of yield stresses (25 ≤ σ_y_ ≤ 40 Pa; Supp. Fig. S12), suggesting that matrix yield stress may also affect downstream differentiation. To further test this idea, we examined the formation of radially-arranged and lumenized neural rosettes. Rosettes are a critically important early indicator of appropriately differentiating MBOs^[43]^, and are thought to be the sites of early diversification in differentiating cell types^[42]^. We therefore analyzed the number and size of these internal structures and found that the number of rosettes consistently increased with matrix yield stress (Fig. 5c, e), but the size of these rosettes peaked at yield stresses of ∼25 Pa (Fig. 5f). This is quite reasonable as rosette size must be dependent upon the ability of the organoid to expand in size to maintaining and enlarge an internal lumen, and higher yield stresses would restrict this expansion. Interestingly, relatively few but large rosettes form in rBM (Supp. Fig. S12), suggesting that tuning yield stress may be critical in recreating some of the mechanical complexity presented in this ubiquitous natural biomaterial. In contrast, increased matrix stiffness alone has been shown only to decrease the number and size of rosettes^[43]^, suggesting that tuning matrix stiffness and yield stress are functionally distinctive parameters in designing biomaterials.

## 3. Discussion

The tunable yield-stress biomaterial developed here acts as a stress-regulating matrix that both supports and limits the generation of locally maintained stresses during tissue growth and morphogenesis. In most ECM, plastic adaptations and remodeling are tightly coupled to local secretory or enzymatic biochemical activity, or to the viscoelastic stress relaxation of the polymer network over time. Hence, while mechanical stresses can be characterized during morphogenesis, it remains challenging to establish a causal link between those stresses and tissue development. The granular gels developed in this work uniquely allow us regulate tissue stresses spatially and autonomously, while minimizing changes in other variables such as stiffness, to clearly define the importance of matrix-mediated mechanical stresses on tissues during development.

We first demonstrate that this material can be tuned over a sufficient yield stress range to limit non-adhesive migration mechanisms of cancer cells. Interestingly, very small changes in yield stress of only ∼15Pa are sufficient to reliably delay observable migration, demonstrating that this biomaterial design can tune yield stress with sufficient resolution to impact biological function, and that biological systems are remarkably sensitive to even small changes in matrix yield stress. MBO developmental studies further support these findings: the yield stress required to achieve maximal changes in morphology and optimal production of large neural rosettes are similarly precise (within a 15 Pa range), and unexpectedly small (σ_y_ ∼ 25 Pa). This value is considerably less than the 100-400 Pa range of accepted measures of yield stress in cell-free Matrigel^[46]^. However, little is known about the yield stress at the organoid-matrix interface, which must be affected by biochemical and enzymatic activity as well as applied mechanical stress. These results highlight the need for technologies to analyze mechanics in situ, as studying cell-free materials do not capture the complex mechanobiological feedback loops that are present even in simple systems; as well as the exquisite sensitivity of biological systems to changes of even 10s of Pascals in stress development.

Broadly, these experiments suggest that yield stress is a critically important mechanical parameter to consider when designing alternatives to natural rBM materials such as Matrigel^[47]^. Our studies demonstrate that a small amount of stress must be maintained for appropriate development, suggesting a targetable metric in designing the mechanical contributors of such scaffolding materials. However, given that controlled uniform yield stresses still do not produce the rosette sizes observed in standard Matrigel, there must be other spatial and temporal mechanical cues required to drive tissue development. Speculatively, perhaps spatial and temporal heterogeneity in yield stress are both required to fully recapitulate MBO morphology. If we conceptualize MBO morphogenesis as a squishy mesh stress ball (graphical abstract), high yield stresses would be required to generate sufficiently large stored internal growth stresses to form buds, after which localized yielding at specific sites would then allow expansion and bud development.

While this material serves as an excellent platform for mechanobiological hypothesis testing, some additional caveats and future directions must be considered. First, the polyacrylamide granules are not adhesive to cells, and hence only resist growth stresses that are normal to the growth surface. Introducing orthogonally-grafted cell-adhesive ligands into these materials would afford the ability to explore shear-generating mechanisms in tissue growth and development, which could lead to further insights into mechanobiological regulators at the tissue-matrix interface; but would also require complex analyses to develop an understanding of the stresses within the tissue. Second, an ideal mechanobiological substrate to study yield stress would keep all other mechanical parameters constant across the entire tunable range. While this is primarily a concern in cell-adhesive materials, and we do successfully establish that the material stiffness is relatively constant over a defined range of yield stresses, strategies to extend this range may be of great value. Given the established importance of matrix stiffness in regulating function, dynamically controlling stiffness in conjunction with yield stress could also produce considerable insights into fundamental mechanobiological relationships between cells and ECM. Third, ideal materials would provide optical clarity to support real-time imaging. As observed in our peripheral shedding demonstration, the refractive indices at the interfaces between granules and media is mismatched and cannot be predicted or compensated for in microscopy imaging or post-processing, making it challenging to perform high-quality imaging live studies. While we extracted the tissues for high-magnification image analysis after fixation, developing refractive index optical matching technologies that are compatible with live imaging requirements could greatly improve the utility of these granular materials as experimental substrates for mechanobiological studies.

## 4. Conclusion

This study introduces a stress-adaptive granular hydrogel platform that enables precise tuning of yield stress while maintaining near-constant stiffness, providing a unique tool to decouple plasticity from other mechanical parameters in 3D culture systems. By leveraging concentration-dependent guest–host supramolecular interactions on polyacrylamide granules, we achieved cohesive matrices with biologically relevant yield stresses ranging from 12 to 370 Pa. These materials regulate growth-induced stresses in vitro, influencing both invasive behavior of breast cancer spheroids and morphogenetic processes in midbrain organoids. Remarkably, subtle changes in yield stress on the order of tens of pascals significantly altered organoid architecture, including neural rosette formation, underscoring the sensitivity of developmental programs to mechanical plasticity. This platform addresses a critical gap in mechanobiology by enabling hypothesis-driven studies of stress-regulated tissue development and disease progression. Beyond fundamental research, these findings suggest yield stress as an important design parameter for next-generation biomaterials in regenerative medicine, disease modeling, and biomanufacturing applications.

## 5. Experimental Section

Unless stated otherwise, all cell culture materials and supplies were purchased from Fisher Scientific (Ottawa, ON) and chemicals from Sigma Aldrich (Oakville, ON).

### 5.1 Preparation of β-cyclodextrin- and adamantane-acrylamides and labels

βCD-AAm and Ad-AAm were synthesized as previously described^[27]^. 6-Amino-βCD was prepared in a sodium bicarbonate solution (0.1 M; pH adjusted to 10 with sodium hydroxide). Acryloyl chloride was added dropwise and left to react for 8 hours. The solution was then evaporated to 40% of the total volume and precipitated in acetone. The precipitate was collected and freeze-dried. The product was purified with an HP-20 polystyrene-divinylbenzene gel using a gradient of water/methanol to yield βCD-AAm. To make Ad-AAm, adamantylamine and triethylamine were dissolved in tetrahydrofuran over ice. Acryloyl chloride was added dropwise and left to react overnight. The precipitate was removed by filtration, and the supernatant was freeze-dried overnight. Using silica gel column chromatography, the product was eluted with hexane/ethyl acetate to yield Ad-AAm. To make Ad-Fluorescein and βCD-Fluorescein for use as labelling agents against the complementary moiety, 6-Amino-βCD or Adamantylamine was dissolved in PBS and NHS-Fluorescein was added in 1:1.25 ratio and allowed to react overnight. The resulting labels were then added in a stoichiometrically excessive volume to the substrates when required.

### 5.2 Polyacrylamide bulk gel preparation

Bulk PAAm gels were created for use as described previously (Table 1)^[48]^. Acrylamide (Aam; Bio-Rad) was combined with bis-acrylamide (Bis; Bio-Rad) in phosphate buffered saline (PBS), along with tetramethylethylenediamine (TEMED). Freshly prepared ammonium persulfate (APS) (Bio-Rad) in PBS was added to the AAm solution to initiate the reaction. The mixture was vortexed, pipetted onto 18 mm cover slips, covered with an additional coverslip, and allowed to gel at room temperature. Gels were transferred directly to a 6-well plate, covered in PBS, and left for 1 hour on a shaker plate to aid in coverslip removal. Coverslips were removed, and gels were washed for 30 minutes twice in fresh PBS. Plates were sealed with Parafilm and stored at 4°C at least overnight to ensure swelling, or until use.

**Table 1.** PAAm gel formulations for 1000 μL of gel.

| Formation<br>(AAm%/Bis% (w/v)) | 3/0.05 | 3/0.1 | 7.5/0.05 |
| --- | --- | --- | --- |
| 40% Acrylamide (μL) | 75 | 75 | 187.5 |
| 2% Bisacrylamide (μL) | 24.5 | 53.5 | 118.0 |
| PBS (μL) | 799.0 | 770.0 | 684.0 |
| TEMED (μL) | 1.5 | 1.5 | 1.5 |
| 1% APS in PBS (μL) | 100 | 100 | 100 |
| Total | 1000 | 1000 | 1000 |

### 5.3 Granular hydrogel fabrication and handling

Host-guest functionalized hydrogel granules were formed via polymerization in an oil-water emulsion. βCD-AAm was added directly to the pre-polymerized aqueous solutions, but due to low aqueous solubility, the Ad-AAm was dissolved in the oil phase, where it reacts at the oil-water interface to form Ad-coated granules. For those experiments requiring labelled granules, either 0.2 μm carboxylate-modified red latex particles (Fluosphere, Life technologies) or fluorescein o-methacrylate (100mg/mL in DMSO; 1 μL/mL of hydrogel prepolymer) were added to the aqueous phase before polymerization. Aqueous prepolymer, APS initiator, and oil-phase solutions (6% v/v of polyglycerol polyricinoleate (Palsgaard) in kerosene) were separately prepared, and filter sterilized (0.22 μm) into rubber-septum capped glass tubes, and oxygen purged via nitrogen bubbling for 20 minutes. The APS was added and mixed into the aqueous prepolymer and then injected into the kerosene solution via syringe. The two-phase emulsion was vortexed and allowed to polymerize while stirring at 1000 RPM for 15 minutes on a stir plate to form hydrogel granules.

To process the fabricated granules, they were allowed to settle by gravity and washed in fresh kerosene three times to remove the surfactant. Kerosene was then displaced by pelleting the granules at 14,000 RCF for two minutes and resuspended in sterile PBS three times. Granules used for cell culture were then treated in 3% antimycotic/antifungal under UV light for 24 hours, before storage in sterile PBS at 4 ºC overnight. When required, granules were individually manipulated using a TRIO-TM MP-245 Micromanipulator System (Sutter Instruments) via 1 mm borosilicate glass capillaries pulled on a Flaming/Brown Micropipette Puller Model P-97 (Sutter) to produce 1-2 μm diameter openings. Holding suction was manually applied via a syringe connected to the micropipette as needed. For bulk handling, granule slurries were transferred via micropipettes with enlarged tips.

### 5.4 Rheological characterization

Mechanical characterization of hydrogels was performed an Anton Paar MCR 302 Modular Compact Rheometer equipped with an 8 mm parallel plate geometry at 37°C. Granular packed-bed hydrogels were formed by mixing Ad and βCD granules in a 1:1 ratio, and either pipetted or vortexed to ensure homogenous mixing, and centrifuged at 14,000 RCF for 2 minutes, before transfer to the rheometer stage. Shear stress, storage modulus (G’), and loss modulus (G”) were assessed by an amplitude sweep from 0.01 to 1000% shear strain. Elastic modulus was determined from the linear region of G’ reading and yield point identified based on a 5% change from the baseline elastic modulus.

### 5.5 Spheroid cell culture

MCF7 and T47D (ATCC) breast cancer cell lines were cultured in Dulbecco’s Modified Eagle’s Medium (DMEM) high glucose supplemented with 10% (v/v) fetal bovine serum (FBS) and 1% (v/v) anti-anti at 37°C, 5% CO_2_. Culture media was changed every two days. At 80% confluency, cells were passaged using Trypsin-EDTA (0.25% v/v) and re-plated in complete growth medium. Aggregates of T47D cell line were formed in arrays of non-adhesive PAAm microwells with 500 μm diameter^[49]^. Briefly, PAAm microwells were formed by polymerizing a 12/2.5 (AAm%/Bis% (w/v)) formulation against a 3D-printed mold and sterilized as previously described. 200 μL of T47D cell suspension (10×10^6^ cells/mL) were allowed to settle into the microwells for 5 minutes before excess media was replaced. Spheroids were allowed to form for 3 days before embedding in granular gel formulations by centrifugation at 100 RCF for two minutes. Alternatively, spheroids were embedded in linearly-elastic, non-degradable TrueGel hydrogels (TRUE9 kit, Sigma) as per the manufacturers’ instructions. Crosslinking strengths were varied to correspond with an elastic moduli of 30-1000 Pa.

### 5.6 Organoid formation and maintenance

The midbrain organoids (MBOs) were generated as previously describe^[45]^. Briefly, AIW002-2 (Neuro Biobank ID: IPSC0063; derived from a 37 yr old male PBMC, via Sendai virus reprogramming; cell line not listed on the ICLAC Register of Misidentified Cell Lines)^[50]^ human-derived induced pluripotent stem cells (hiPSCs)^[51]^ were cultured to 60-70% confluency, dissociated with Accutase, and pelleted at 200 RCF for 3 minutes. Cell pellets were resuspended in neural induction media (Supporting Table S2) and seeded into U-bottom ultra-low adhesion 96-well plates (200 μL per well at 5×10^4^ cells/mL). Plates were centrifuged at 200 RCF for 10 minutes and incubated at 37°C and 5% CO_2_. After 2 days, ROCK inhibitor was removed from the neural induction culture media formulation. On day 4, the media was exchanged for midbrain patterning formulations (Supporting Table S3).

On day 7, organoids were embedded in either Growth Factor Reduced (GFR) Matrigel; or in 50 μL of the yield-stress tunable granular gel formulations. Organoids that were cultured in packed granular gel were treated with 1.5% GFR-Matrigel added to the neural induction media on day 7 (Supporting Table S4). For these long-term experiments, round-bottom wells (2mm diameter, 3mm depth) were replica-molded in polyacrylamide as previously described and sequentially seeded with (1) 25 μL of each pre-mixed packed-gel formulation (2) MBOs and (3) an additional 25 μL of granules. Each layer was centrifuged at 75 RCF for 8 minutes; to uniformly embed the MBOs. Organoids were then cultured in final differentiation media (Supporting Table S5) for the remainder of the experimental time course.

### 5.7 Fluorescent labelling and tissue clearing

To assess viability, spheroids and organoids were stained with Calcein AM (2 μM) and Ethidium Homodimer-1 (4 μM; Life Technologies) to fluorescently label live and dead cells for 30 minutes, and counterstained with Hoechst 33342 (2 μL/mL in PBS). For immunofluorescent labels, Matrigel was first removed using Cell Recovery Solution (Corning), and tissues were fixed in 4% paraformaldehyde overnight at 4°C. Organoids were then cleared using a previously established protocol^[45]^ in CUBIC reagent on a shaker at 80 RPM and 37°C for 72h (R1, lipid removal); and permeabilized and blocked (0.5% Triton X-100 + 5% donkey serum in PBS) for one hour. Primary antibodies against microtubule-associated protein 2 (MAP2; chicken polyclonal, EnCor Biotechnology CPCA-MAP2), tyrosine hydroxylase (TH; rabbit polyclonal, Pel Freez P40101-150), and neural stem cell (Nestin; mouse polyclonal, DSHB) were diluted by 1:200 in blocking solution, and incubated with the organoids for 72 hours at 37 °C. After thoroughly rinsing the tissues in PBS, secondary fluorescent antibodies and Hoechst 33342 were applied for 72 hours. Finally, organoids were cleared with CUBIC reagent 2 (R2, for refractive index adjustment), and imaged within 24 hours.

### 5.8 Imaging and microscopy

Routine live, brightfield and fluorescent imaging was performed on an EVOS M7000 Cell Imaging System (Fisher). For quantitative analyses, an IX73 Olympus Microscope with ORCA Flash 4.0LT camera was used with a 20x objective. Confocal microscopy was performed using an Opera Phenix Plus High-Content Screening System (Perkin-Elmer; 20X-water objective).

### 5.9 Image Analysis

Size characteristics of granule populations were calculated using standard approaches ^[52]^, for which the diameters of granules were measured manually in FIJI (NIH) ^[53]^. Polydispersity indices were calculated using the population mean and standard deviation as in Equation 1.

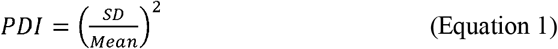

Data collection for assays requiring fluorescent intensity quantification were performed using identical imaging conditions, with integrated density measurements collected in FIJI, after subtraction of integrated background intensity.

Aggregate invasion was described by measuring the aggregate circularity over time, by manually drawing an outline around each aggregate, and calculating circularity^[53]^ as in Equation 2.

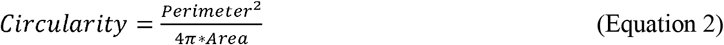

### 5.9 Statistical analysis

Statistical analysis was performed in GraphPad Prism 9.1.2 (GraphPad Software, San Diego, California USA). The normality of each dataset was determined using a Shapiro-Wilk test. Equivalence of variance was assessed using a Brown-Forsythe test. Sphericity was not assumed, and a Geisser-Greenhouse correction was applied. For normally distributed data with equal variances, one-way ANOVAs were applied with Tukey post-hoc tests for multiple comparisons. Repeated measures for one-way ANOVA tests were also used for sequentially collected data. Conditions that did not pass normality were assessed using a Friedman test followed by Dunn’s multiple comparisons. The significance level alpha was set based on the variability of the control conditions (alpha = 0.05 by default, but alpha = 0.001 for invasive growth assays).

### 5.10 Use of AI-based software

Microsoft Copilot was used to generate the schematic in the graphical abstract.

## Supporting Information

Supporting Information is available from the Wiley Online Library or from the authors.

## Acknowledgements

We thank Dr. Wontae Lee, Pooja Patel and Nicholas Stylianesis for initial work on early versions of this project; Dr. Alejandro Forigua Coronado, Christina Boghdady and Benjamin Campbell for helpful discussions; and Taylor Goldsmith and Wolfgang Reintsch for training and insights in stem cell culture and microscopy respectively.

## Data availability statement

Source data is maintained in an Open Science Framework repository that can be accessed at: https://osf.io/cb4gv/overview?view_only=19bfe15fecf34d16b1571571b9dc1ac0

## Ethics approval statement

Approved with McGill REB under Study Number A03-M19-22A / eRAP 22-03-027.

## References

[1] A. Saraswathibhatla, D. Indana, O. Chaudhuri, Nat. Rev. Mol. Cell Biol. 2023, 24, 495.

[2] L. Yang, P. Jiang, J. B. Stein, Y. Hou, C. Zhou, H. Kang, K.-B. Lee, Adv. Sci. n.d., n/a, e16992.

[3] D. Baruffaldi, G. Palmara, C. Pirri, F. Frascella, ACS Appl. Bio Mater. 2021, 4, 2233.

[4] W. Fan, K. Adebowale, L. Váncza, Y. Li, M. F. Rabbi, K. Kunimoto, D. Chen, G. Mozes, D. K.-C. Chiu, Y. Li, J. Tao, Y. Wei, N. Adeniji, R. L. Brunsing, R. Dhanasekaran, A. Singhi, D. Geller, S. H. Lo, L. Hodgson, E. G. Engleman, G. W. Charville, V. Charu, S. P. Monga, T. Kim, R. G. Wells, O. Chaudhuri, N. J. Török, Nature 2024, 626, 635.

[5] N. Kalashnikov, Soft Matter 2023.

[6] X. Dai, D. Wu, K. Xu, P. Ming, S. Cao, L. Yu, ACS Appl. Mater. Interfaces 2025, 17, 8751.

[7] G. S. Major, H. Joukhdar, Y. S. Choi, J. Rnjak-Kovacina, S. G. Wise, L. A. Ju, T. R. Cox, C. Xu, G. C. Yeo, J. L. Young, Cell Rep. Phys. Sci. 2025, 6.

[8] H. T. Ong, M. Sriram, H. H. Susapto, Y. Li, Y. Jiang, N. H. Voelcker, J. L. Young, A. W. Holle, R. Elnathan, Adv. Mater. 2025, 37, 2501640.

[9] M. Lucariello, M. L. Valicenti, S. Giannoni, L. Donati, I. Armentano, F. Morena, S. Martino, Biomolecules 2025, 15, 848.

[10] R. Curvello, V. Kast, P. Ordóñez-Morán, A. Mata, D. Loessner, Nat. Rev. Mater. 2023, 8, 314.

[11] A. Garatikar, A. Raaza, P. K. R., S. R., M. Nedunchezhiyan, P. Prabhakar, Regen. Eng. Transl. Med. 2025, DOI 10.1007/s40883-025-00479-w.

[12] C. Ort, W. Lee, N. Kalashnikov, C. Moraes, Expert Opin. Drug Discov. 2021, 16, 159.

[13] J. M. Grolman, P. Weinand, D. J. Mooney, Proc. Natl. Acad. Sci. 2020, 117, 25999.

[14] S. R. Naganathan, Biochem. Soc. Trans. 2024, 52, 987.

[15] N. D. Leipzig, M. S. Shoichet, Biomaterials 2009, 30, 6867.

[16] J. Lantoine, T. Grevesse, A. Villers, G. Delhaye, C. Mestdagh, M. Versaevel, D. Mohammed, C. Bruyère, L. Alaimo, S. P. Lacour, L. Ris, S. Gabriele, Biomaterials 2016, 89, 14.

[17] M. Matejčić, X. Trepat, Trends Cell Biol. 2023, 33, 95.

[18] M. Tang, S. K. Tiwari, K. Agrawal, M. Tan, J. Dang, T. Tam, J. Tian, X. Wan, J. Schimelman, S. You, Q. Xia, T. M. Rana, S. Chen, Small 2021, 17, 2006050.

[19] M. Wang, Y. Yang, L. Han, S. Han, N. Liu, F. Xu, F. Li, Biochem. Biophys. Res. Commun. 2020, 528, 459.

[20] C.-M. Boghdady, N. Kalashnikov, S. Mok, APL Bioeng. 2021.

[21] B. Buchmann, P. Fernández, A. R. Bausch, Biophys. Rev. 2021, 2, 021401.

[22] J. Kim, J. Feng, C. A. R. Jones, X. Mao, L. M. Sander, H. Levine, B. Sun, Nat. Commun. 2017, 8, 842.

[23] B. Buchmann, L. K. Engelbrecht, P. Fernandez, F. P. Hutterer, M. K. Raich, C. H. Scheel, A. R. Bausch, Nat. Commun. 2021, 12, 2759.

[24] Y. Jia, Y. Wang, L. Niu, H. Zhang, J. Tian, D. Gao, X. Zhang, T. J. Lu, J. Qian, G. Huang, F. Xu, Adv. Healthc. Mater. 2021, 10, e2001856.

[25] O. Chaudhuri, J. Cooper-White, P. A. Janmey, D. J. Mooney, V. B. Shenoy, Nature 2020, 584, 535.

[26] A. E. Tonelli, G. Narayanan, A. Gurarslan, Polymers 2018, 10, 911.

[27] T. Kakuta, Y. Takashima, A. Harada, Macromolecules 2013, 46, 4575.

[28] T. H. Qazi, V. G. Muir, J. A. Burdick, ACS Biomater. Sci. Eng. 2022, 8, 1427.

[29] Y. Jin, A. Compaan, T. Bhattacharjee, Y. Huang, Biofabrication 2016, 8, 025016.

[30] V. G. Muir, T. H. Qazi, S. Weintraub, B. O. Torres Maldonado, P. E. Arratia, J. A. Burdick, Small 2022, 18, 2201115.

[31] Y. Jin, A. Compaan, T. Bhattacharjee, Y. Huang, Biofabrication 2016, 8, 025016.

[32] V. R. Feig, S. Santhanam, K. W. McConnell, K. Liu, M. Azadian, L. G. Brunel, Z. Huang, H. Tran, P. M. George, Z. Bao, Adv. Mater. Technol. 2021, 6, 2100162.

[33] R. J. Pelham, Y. Wang, Proc. Natl. Acad. Sci. U. S. A. 1997, 94, 13661.

[34] L. Zhao, S. Mok, C. Moraes, Biofabrication 2019.

[35] A. J. Engler, S. Sen, H. L. Sweeney, D. E. Discher, Cell 2006, 126, 677.

[36] W. Lee, N. Kalashnikov, S. Mok, R. Halaoui, E. Kuzmin, A. J. Putnam, S. Takayama, M. Park, L. McCaffrey, R. Zhao, R. L. Leask, C. Moraes, Nat. Commun. 2019, 10, 144.

[37] W. Lee, C.-M. Boghdady, V. Lelarge, R. L. Leask, L. McCaffrey, C. Moraes, Biomaterials 2023, 296, 122073.

[38] B. E. Campbell, K. Zhang, A. Shi, S. Rostami, D. Pioche-Lee, C. Li, A. Leblond, A. Forigua, C.-M. Boghdady, C. Moraes, S. C. Lesher-Pérez, Appl. Bio Mater. 2025.

[39] K. M. Young, C. A. Reinhart-King, Curr. Opin. Cell Biol. 2023, 83, 102208.

[40] A. B. Behrooz, S. Shojaei, Biochim. Biophys. Acta BBA - Mol. Basis Dis. 2024, 1870, 167332.

[41] O. Piwocka, K. Sterzyńska, A. Malińska, W. M. Suchorska, K. Kulcenty, Sci. Rep. 2025, 15, 27449.

[42] M. A. Lancaster, M. Renner, C.-A. Martin, D. Wenzel, L. S. Bicknell, M. E. Hurles, T. Homfray, J. M. Penninger, A. P. Jackson, J. A. Knoblich, Nature 2013, 501, 373.

[43] C. Cassel de Camps, S. Aslani, N. Stylianesis, H. Nami, N.-V. Mohamed, T. M. Durcan, C. Moraes, ACS Appl. Bio Mater. 2022, 5, 214.

[44] S. Nam, J. Lee, D. G. Brownfield, O. Chaudhuri, Biophys. J. 2016, 111, 2296.

[45] N.-V. Mohamed, M. Mathur, R. V. da Silva, R. A. Thomas, P. Lepine, L. K. Beitel, E. A. Fon, T. M. Durcan, MNI Open Res. 2022, 3, 1.

[46] J. Reed, W. J. Walczak, O. N. Petzold, J. K. Gimzewski, Langmuir ACS J. Surf. Colloids 2009, 25, 36.

[47] E. A. Aisenbrey, W. L. Murphy, Nat. Rev. Mater. 2020, 5, 539.

[48] J. R. Tse, A. J. Engler, Curr. Protoc. Cell Biol. 2010, 47, 10.16.1.

[49] L. Zhao, S. Mok, C. Moraes, 2019.

[50] “CBIGi001-A · Cell Line · hPSCreg,” can be found under https://hpscreg.eu/cell-line/CBIGi001-A, n.d.

[51] C. X.-Q. Chen, N. Abdian, G. Maussion, R. A. Thomas, I. Demirova, E. Cai, M. Tabatabaei, L. K. Beitel, J. Karamchandani, E. A. Fon, T. M. Durcan, Methods Protoc. 2021, 4, 50.

[52] N. Raval, R. Maheshwari, D. Kalyane, S. R. Youngren-Ortiz, M. B. Chougule, R. K. Tekade, in Basic Fundam. Drug Deliv. (Ed.: R. K. Tekade), Academic Press, 2019, pp. 369–400.

[53] J. Schindelin, I. Arganda-Carreras, E. Frise, V. Kaynig, M. Longair, T. Pietzsch, S. Preibisch, C. Rueden, S. Saalfeld, B. Schmid, J.-Y. Tinevez, D. J. White, V. Hartenstein, K. Eliceiri, P. Tomancak, A. Cardona, Nat. Methods 2012, 9, 676.

