## Supplementary material for "Stress-adaptive biomaterials with tunable yielding architectures regulate organoid morphogenesis": n/a

1. **Supporting Movies**

**Supporting Movie SM1** – non-attachment between unfunctionalized beads on glass capillaries. Stills in Supplemental Figure S3

**Supporting Movie SM2** – attachment between functionalized beads on glass capillaries. Stills in Manuscript Figure 3c.

**Supporting Movie SM3** – mixing of beads to form solid-like material. Still in manuscript Figure 3e.

2. Supporting Figures

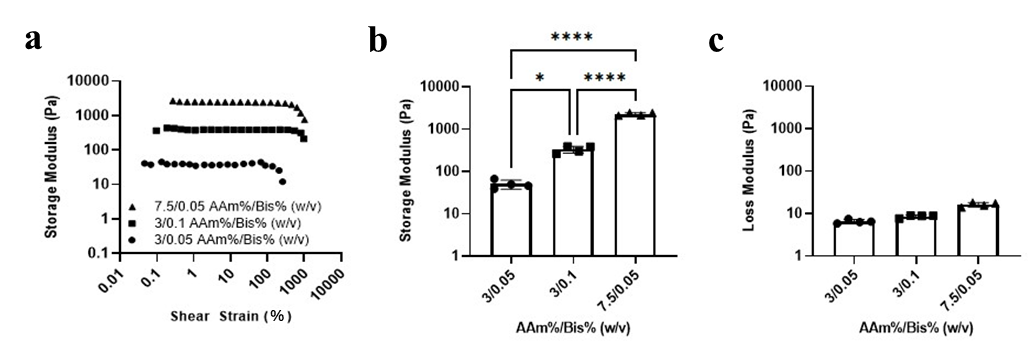

**Supporting Figure S1. Stiffness of hydrogel formulations** **used to make granules**. **(a)** shear stress amplitude sweeps of bulk polyacrylamide hydrogel formulations on a rheometer, to obtain **(b)** storage and **(c)** loss moduli for the gels. Since polyacrylamide granules formed in an oil-water emulsion maintain similar mechanical properties to gels fabricated in bulk^[1]^, these storage modulus values are a reasonable approximation for the moduli of individual granules. Gel stiffness was tunable via gel formulation within a physiologically relevant range (means G = 0.05, 0.2, and 2.5 kPa, with negligible loss moduli.) Data presented as mean ± standard deviation, n = 4-6, * p < 0.05, ** p < 0.01, **** p < 0.0001 by one-way ANOVA with Tukey posthoc comparisons.

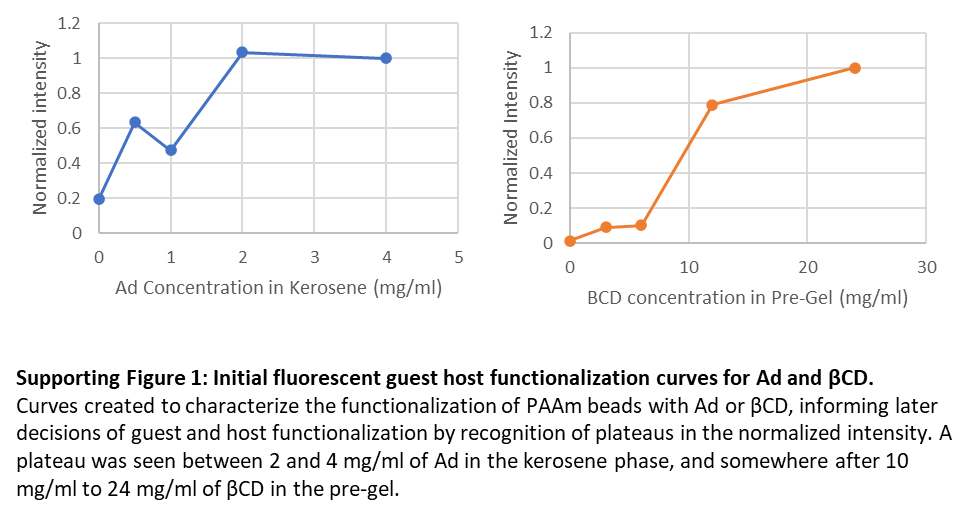

**Supporting Figure S2: Degree of functionalization for Ad and βCD on hydrogel granules.** Was used to identify relative maxima of functionalization concentration of both Adamentane and Beta-Cyclodextrin. Plateaus were observed between 2 and 4 mg/mL of Ad in the kerosene phase, and between 10 and 24 mg/mL of βCD in the pre-gel.

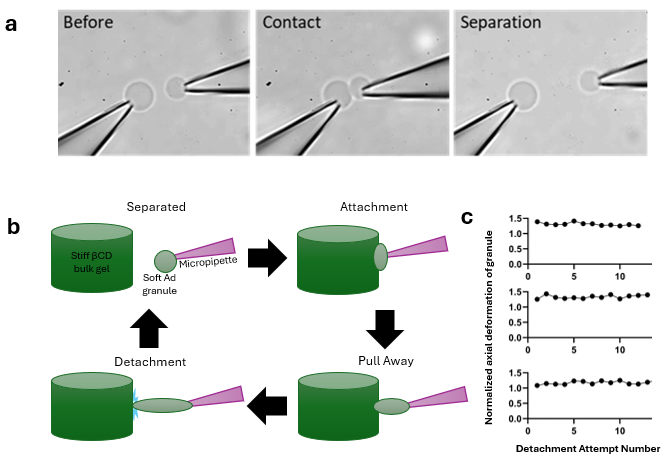

**Supporting Figure S3: Micromanipulation of PAAm granules**. **(a)** Contact between unfunctionalized PAAm granules demonstrated no binding, in contrast with contact between guest- and host-functionalized granules (shown in manuscript Figure 3c). **(b)** Repeated binding assay in which a granule is micromanipulated against a complementarily functionalized gel surface and pulled away. **(c)** Granule deformation for repeated cycles of attachment/detachment, demonstrating that repeated binding occurs over multiple attachment/detachment events. Data is shown for three individual beads undergoing > 10 attachment/detachment cycles.

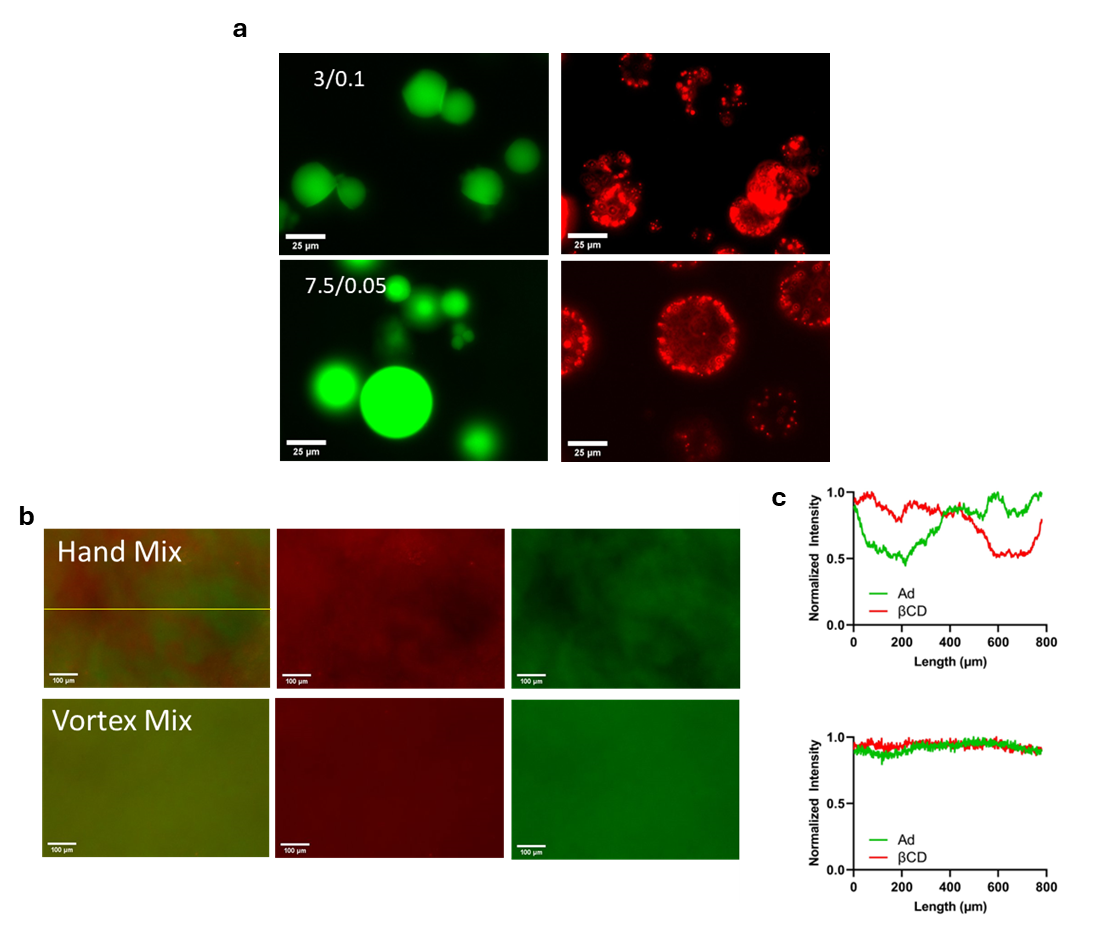

**Supporting Figure S4:** **Behavior of guest and host functionalized beads when mixed. (a)** Ad-functionalized (guest; labelled in green) and βCD-functionalized (host; labelled in red) granules of 3/0.1 and 7.5/0.05 AAm/Bis (w/v%) were mixed. Soft gels deformed more during binding (3/0.1, top) when compared to stiffer formulations (7.5/0.05, bottom). Note: figures display the separated fluorescent channels shown merged in manuscript Figure 3a, scale bar 25 µm. **(b)** Samples were mixed by vigorous pipetting (top) or vortexed and centrifuged (bottom) to form packed granules. Scale bar 100 µm. **(c)** Normalized fluorescent intensity of hand-pipetted (top) and vortex-mixed (bottom) packed granular gels.

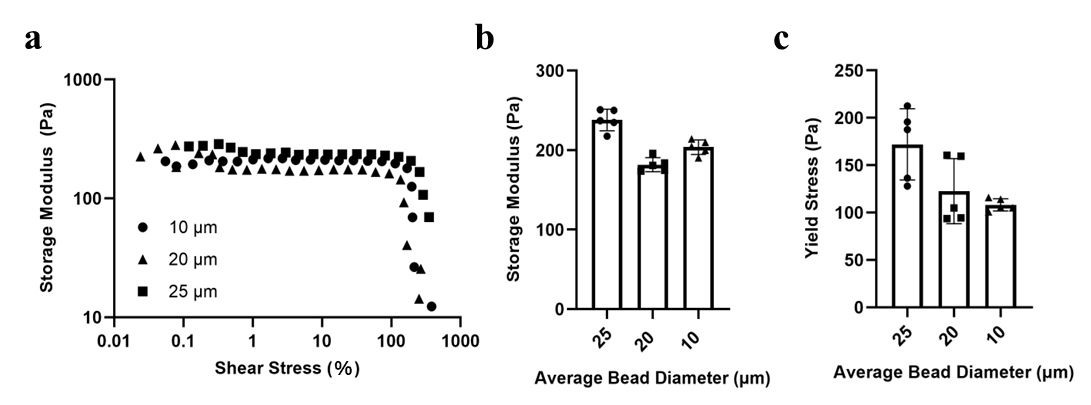

**Supporting Figure S5: Rheology data of packed granular gels based on bead size. (a)** Representative rheometry amplitude sweeps of packed granular gels made of unfunctionalized PAAm granules of differing sizes. **(b)** Storage modulus of packed granular gels made of unfunctionalized PAAm granules of different sizes. **(c)** Yield stress of packed granular gels made of unfunctionalized PAAm granules of different sizes.

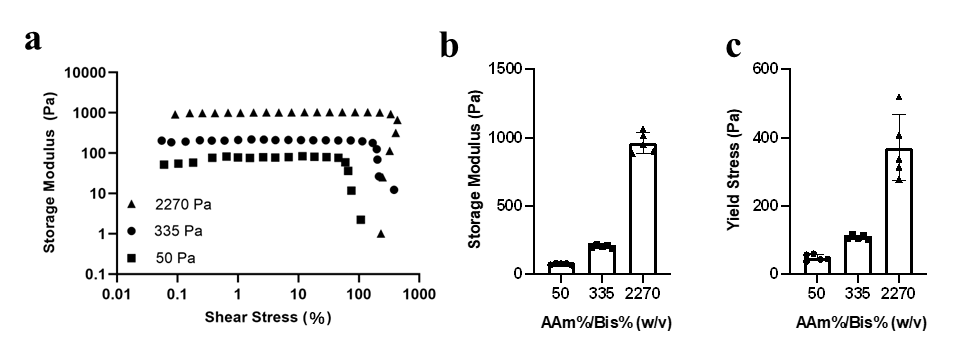

**Supporting Figure S6: Rheology data of packed granular gels based on variations in the stiffness and polymer content of individual granules (a)** Representative amplitude sweeps, **(b)** Storage modulus, and **(c)** Yield stress of packed granular gels made of unfunctionalized PAAm granules with varied base monomer concentrations (details in Supplemental Table S1A).

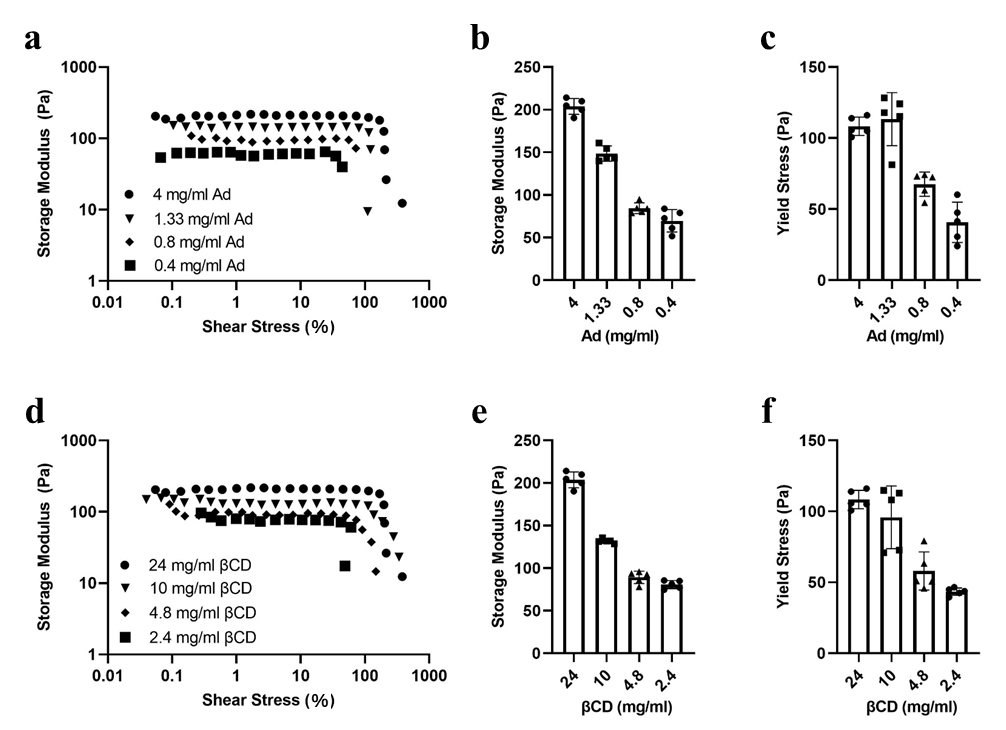

**Supporting Figure S7: Rheological characterization of granular packed gels.** Packed granular guest-host gels were fabricated and mechanically analyzed by shear rheometry. After setting the base granule fabrication parameters (120s vortex time, 3/0.1 AAm%/Bis% (w/v)) we obtained **(a)** amplitude sweeps, **(b)** storage moduli and **(c)** yield stress of granular packed gels with varied Ad concentration against 24 mg/mL of βCD-functionalized granules. **(d)** Amplitude sweeps, **(e)** storage moduli and **(f)** yield stress of granular packed gels with varying βCD concentration against 4 mg/mL of Ad-functionalized granules.

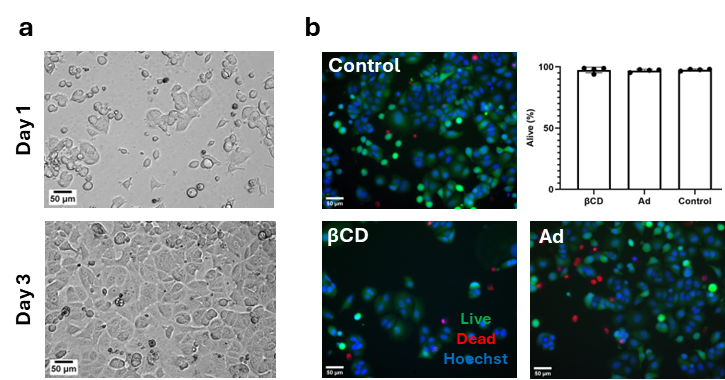

**Supporting Figure S8: Live dead test of functionalized PAAm granular hydrogels with MCF7 breast cancer cells. (a)** day 1 and day 3 of cells in culture. **(b)** cell viability on day 3 for βCD, Ad, and unfunctionalized beads.

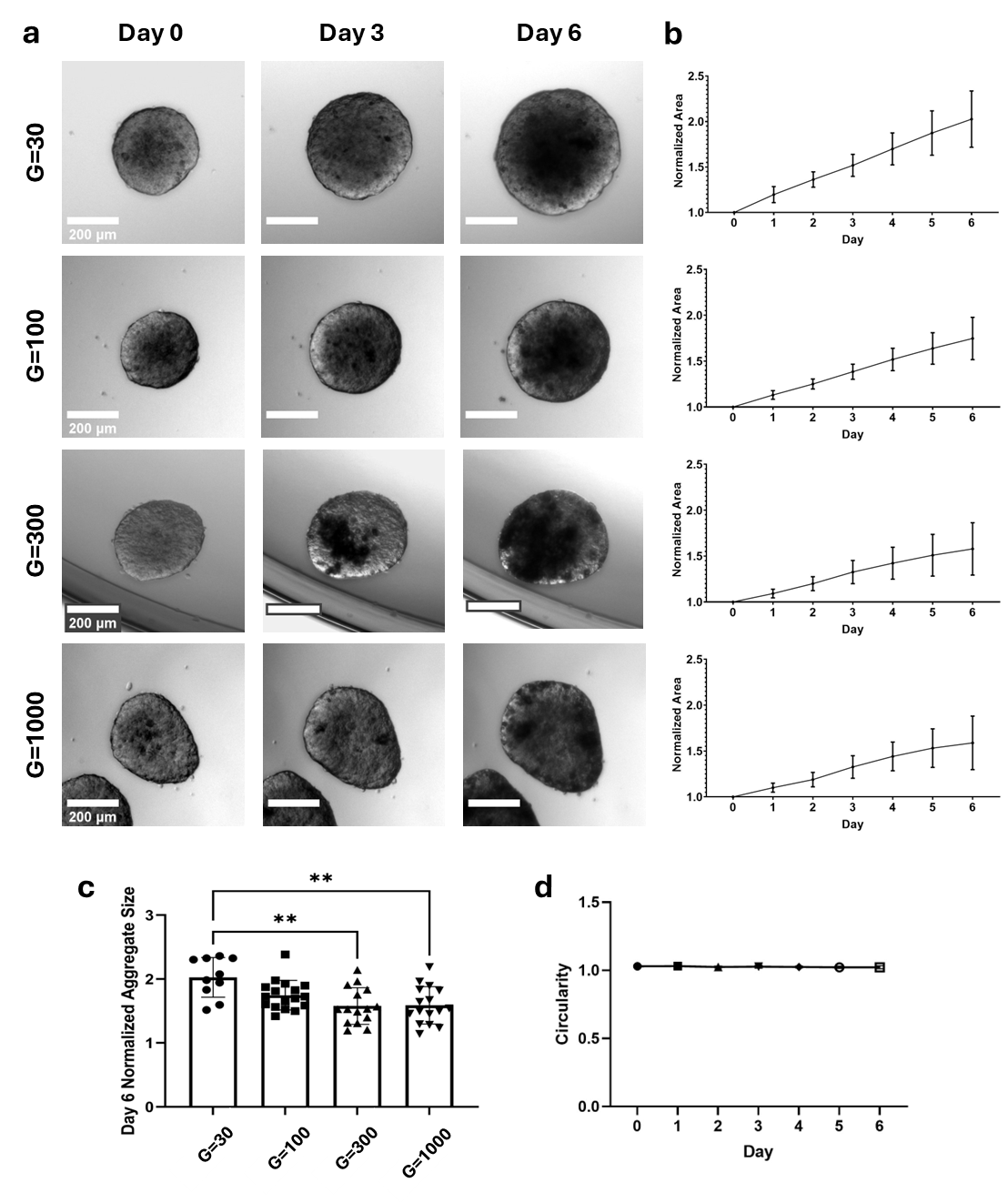

**Supporting Figure S9: Growth of T47D aggregates in non-degradable hydrogel materials with tunable elastic modulus. (a)** Aggregates were embedded in TrueGel (dextran, PVA, and PEG-based commercially available gel system). **(b, c)** Areal growth rates were quantified over 6 days in culture, and **(c)** showed significant differences in normalized size only when the matrix increased in stiffness (G = storage modulus) by 3-10 fold (Data presented as mean ± standard deviation, ** p < 0.01 by ANOVA with Tukey post-hoc comparisons; n = 4-6). **(d)** Aggregate circularity did not change over the culture time.

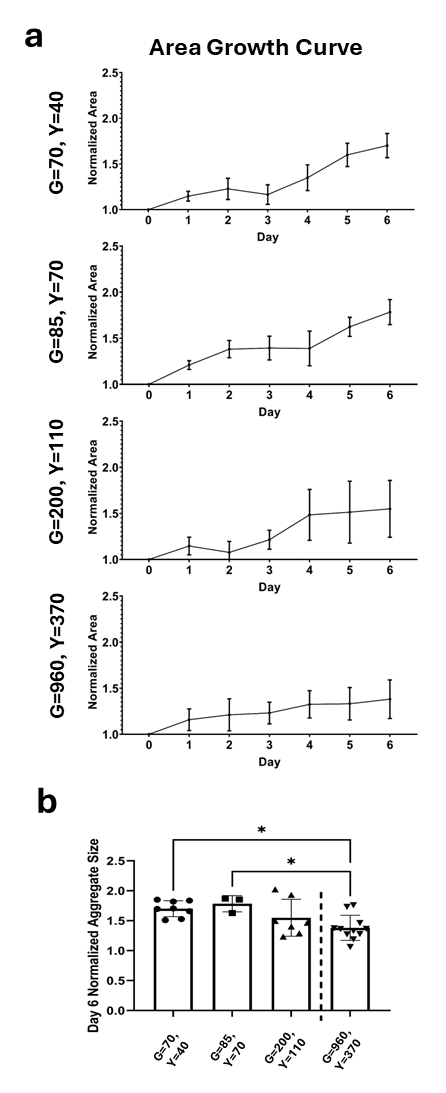

**Supporting Figure S10: Growth of T47D aggregates in elastic TrueGel hydrogels and plastic granular hydrogels. (a)** normalized area of aggregates in TrueGel platforms of different stiffnesses. **(b)** normalized area of aggregates in granular hydrogels of different stiffnesses and yield points. **(c)** Comparison of day 6 aggregate sizes in TrueGel of differing stiffnesses. **(d)** Comparison of day 6 aggregate sizes in packed granular hydrogels of differing stiffness and yield points. (Data presented as mean ± standard deviation, * p < 0.05 by ANOVA with Tukey post-hoc comparisons; n = 4-6) G = elastic modulus (Pa), Y = yield point (Pa).

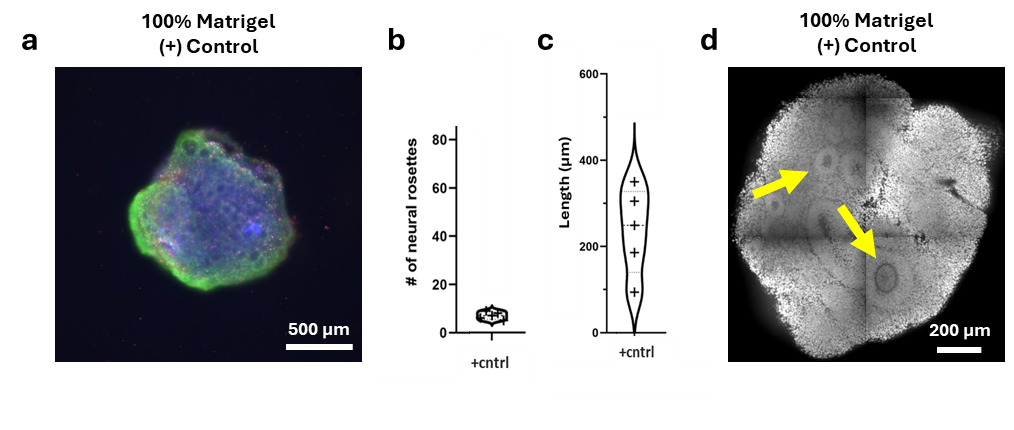

**Supporting Figure S11: Midbrain organoid embedded in 100% Matrigel for 2 weeks. a)** Viability of midbrain organoid embedded in 100% Matrigel (positive control). **b)** Number of neural rosettes in midbrain organoid embedded in 100% Matrigel (positive control). **c)** Average rosette trace length of midbrain organoid embedded in 100% Matrigel (positive control). **d)** Representative whole-mount-stained midbrain organoid embedded in 100% Matrigel (positive control) for nuclei. Yellow arrows point at neural rosettes in the organoids. (All data presented as mean ± standard deviation, n = 4-5)

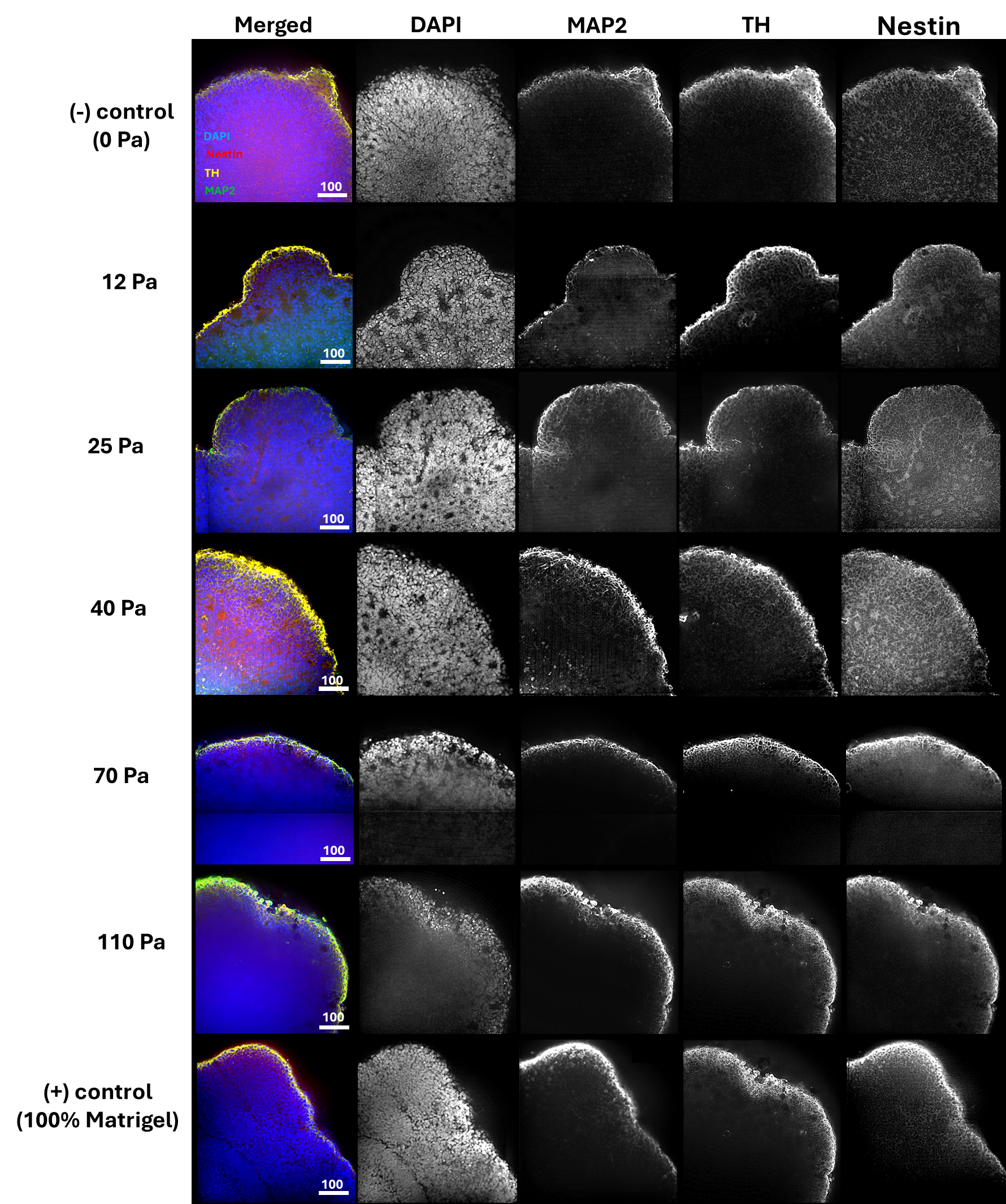

**Supporting Figure S12. Representative whole-****mount stained midbrain organoids 2 weeks post embedding in packed gel.** Microtubule-associated protein 2 (MAP2) indicating the presence of mature neurons, tyrosine hydroxylase (TH) indicating the presence of dopaminergic neurons, neural stem/progenitor cells (Nestin), and DAPI showing cell nuclei. Scale bar = 200 µm

**Supporting Table S1A.** Fabrication parameters used for packed granular gels in T47D aggregate cultures.

| **AAm%/Bis% [w/v]** | **Vortex Time**  **[s]** | **Ad Concentration**  **[mg/mL]** | **βCD Concentration [mg/mL]** | **Storgae Modulus**  **[Pa]** | **Yield Stress**  **[Pa]** |
| --- | --- | --- | --- | --- | --- |
| 3/0.1 | 120 | 0.4 | 24 | 70 ± 12 | 41 ± 18 |
| 3/0.1 | 120 | 0.8 | 24 | 85 ± 8 | 68 ± 10 |
| 3/0.1 | 120 | 4 | 24 | 204 ± 12 | 108 ± 8 |
| 7.5/0.05 | 120 | 4 | 24 | 962 ± 89 | 371 ± 121 |

**Supporting Table S1B.** Fabrication parameters used for packed granular gels in midbrain organoid cultures.

| **AAm%/Bis% [w/v]** | **Vortex Time**  **[s]** | **Ad Concentration**  **[mg/mL]** | **βCD Concentration [mg/mL]** | **Storage**  **Modulus**  **[Pa]** | **Yield Stress**  **[Pa]** |
| --- | --- | --- | --- | --- | --- |
| 3/0.1 | 120 | 0.13 | 24 | 51.9 ± 7.5 | 12.2 ± 2.8 |
| 3/0.1 | 120 | 0.27 | 24 | 57.25 ± 6.4 | 24.9 ± 5.3 |
| 3/0.1 | 120 | 0.4 | 24 | 70 ± 12 | 41 ± 18 |
| 3/0.1 | 120 | 0.8 | 24 | 85 ± 8 | 68 ± 10 |

**Supporting Table S2**. Composition of neuronal induction medium.

| Final Concentration | Recipe (100 ml) |
| --- | --- |
| DMEM/F-12 + Neurobasal (1:1) | 50 mL + 50 mL |
| 1:100 N2 | 1 mL |
| 1:50 B27 without vitamin A | 2 mL |
| 1% GlutaMAXTM-I | 1 mL |
| 1% MEM-NEAA | 1 mL |
| 1:100 2-mercaptoethanol | 35 µL |
| 1 μg/mL Heparin | 100 µL |
| 10 μM SB431542 | 100 µL |
| 200 ng/mL Noggin | 100 µL |
| 0.8 μM CHIR99021 | 27 µL |
| 10 μM ROCK inhibitor | 100 µL |

**Supporting Table S3**. Composition of midbrain patterning medium.

| Final Concentration | Recipe (100 ml) |
| --- | --- |
| DMEM/F-12 + Neurobasal (1:1) | 50 mL + 50 mL |
| 1:100 N2 | 1 mL |
| 1:50 B27 without vitamin A | 2 mL |
| 1% GlutaMAXTM-I | 1 mL |
| 1% MEM-NEAA | 1 mL |
| 1:100 2-mercaptoethanol | 35 µL |
| 1 μg/mL Heparin | 100 µL |
| 10 μM SB431542 | 100 µL |
| 200 ng/mL Noggin | 100 µL |
| 0.8 μM CHIR99021 | 27 µL |
| 200ng/mL SHH | 50µL |
| 100 ng/ mL FGF8 | 100 µL |

**Supporting Table S4**. Composition of tissue induction medium.

| Final Concentration | Recipe (100 ml) |
| --- | --- |
| Neurobasal | 100 mL |
| 1:100 N2 | 1 mL |
| 1:50 B27 without vitamin A | 2 mL |
| 1% GlutaMAXTM-I | 1 mL |
| 1% MEM-NEAA | 1 mL |
| 1:100 2-mercaptoethanol | 35 µL |
| 2,5 μg/mL insulin | 25 µL |
| 200 ng/mL laminin | 17 µL |
| 200 ng/mL SHH | 50 µL |
| 100 ng/ mL FGF8 | 100 µL |
| Pen/Strep | 100 µL |

**Supporting Table S5**. Composition of final differentiation medium.

| Final Concentration | Recipe (100 ml) |
| --- | --- |
| Neurobasal | 100 mL |
| 1:100 N2 | 1 mL |
| 1:50 B27 without vitamin A | 2 mL |
| 1% GlutaMAXTM-I | 1 mL |
| 1% MEM-NEAA | 1 mL |
| 1:100 2-mercaptoethanol | 35 µL |
| 10 ng/mL BDNF | 50 µL |
| 10 ng/mL GDNF | 50 µL |
| 100 μM ascorbic acid | 50 µL |
| 125 μM db-cAMP | 50 µL |
| Pen/Strep | 100 µL |
